# Variability of the urinary and blood steroid profiles in healthy women

**DOI:** 10.1101/2022.10.13.511896

**Authors:** Basile Moreillon, Olivier Salamin, Bastien Krumm, Loredana Iannella, Francesco Molaioni, Tiia Kuuranne, Raul Nicoli, Jonas J. Saugy, Francesco Botrè, Raphael Faiss

**Affiliations:** Research and Expertise in anti-Doping Sciences (REDs), Institute of Sport Sciences, University of Lausanne, Lausanne, Switzerland; Swiss Laboratory for Doping Analyses, University Center of Legal Medicine, Lausanne and Geneva, Lausanne University Hospital and University of Lausanne, Switzerland; Laboratorio Antidoping, Federazione Medico Sportiva Italiana, Rome, Italy

**Keywords:** urine, blood, serum, variability, anti-doping, contraception, women

## Abstract

The steroidal module of the Athlete Biological Passport (ABP) targets the use of exogenous androgenous anabolic steroids (EAAS) in elite sport by monitoring urinary steroid profiles.

Urine and blood samples were collected weekly during two consecutive OCP cycles (8 weeks) in 15 physically active women to investigate the low urinary steroid concentrations and putative confounding effect of OCP.

In urine, testosterone (T) and/or epitestosterone (E) were below the limit of quantification of 1 ng/mL in 62% of the samples. Biomarkers’ variability ranged between 31% and 41%, with a significantly lesser variability for ratios (with the exception of T/E (41%)): 20% for androsterone/etiocholanolone (p < 0.001) and 25% for 5α-androstane-3α,17β-diol/5ß-androstane-3α,17β-diol (p < 0.001).

In serum, variability for testosterone (T; 24%), androstenedione (A4; 23%), dihydrotestosterone (DHT; 19%) and T/A4 (16%) was significantly lower than urinary biomarkers (p < 0.001). Urinary A/Etio increased by > 18% after the first two weeks (p < 0.05) following blood loss. In contrast, T (0.98 nmol/L during the first week), and T/A4 (0.34 the first week) decreased significantly by more than 25% and 17% (p<0.05), respectively in the following weeks.

Our results outline steroidal variations during the OCP cycle highlighting exogenous hormonal preparations as confounder for steroid concentrations in blood. Low steroid levels in urine samples have a clear detrimental impact on the subsequent interpretation of steroidal variations for the ABP. With a greater analytical sensitivity and lesser variability for steroids in serum vs. urine in healthy active women, serum represents a complementary matrix to urine in the ABP steroidal module.

## Introduction

Anabolic androgenic steroids (AAS) are common drugs used as performance enhancing substances. They are intended to increase lean body mass, muscle mass and strength but also to improve recovery when administered in lower doses.^1,2^ In order to detect AAS abuse, by xenobiotics similar to endogenous AAS in particular, the World Anti-Doping Agency (WADA) implemented in 2014 the steroidal module of the Athlete Biological Passport (ABP), which longitudinally monitors a number of testosterone-related substances in urine.^3,4^ The ABP is an adaptive model, which tracks a number of parameters to detect deviations from a normal physiological condition of an individual. Based on population means and previous individual samples, the model computes upper and lower thresholds and may signal suspicious alterations in case of values outside those calculated limits. The following biomarkers are considered in the steroidal module: testosterone (T), androsterone (A), etiocholanolone (Etio), 5α-androstane-3α, 17β-diol (5αAdiol), 5β-androstane-3α,17β-diol (5βAdiol) and epitestosterone (E). These urinary concentrations are then combined within five ratios (T/E, A/Etio, 5αAdiol/E, 5αAdiol/5βAdiol, and A/T) on which the adaptive model is applied.^5^

Large inter- and intra-individual variations among those biomarkers are reported in the literature, especially in female which have far inferior reference ranges than male, particularly for the primary marker T/E.^6–10^ Furthermore, a number of studies have shown that the menstrual cycle has a great influence on steroid biosynthesis and that it induces high intra-individual variations among female populations for whom steroid concentrations are generally low and close to the detection limits.^6^ Due to the variations of E in function of the menstrual phase, E and its associated ratios are particularly subject to high fluctuations, which impairs the detection capacity of the T/E ratio.^11,12^ Additionnaly, hormonal contraceptives may also impact the steroidal metabolism by decreasing the urinary excretion of all ABP metabolites with E and consequently T/E and 5αAdiol/E being the most affected biomarkers.^13–15^ Altogether, these features may confuse the interpretation of female individual profiles of the steroidal module of the ABP, especially for the E-dependent ratios.^12,16^

Currently, urine is the matrix of choice for the application of the steroidal module of the ABP. To complement the drawbacks related to the use of urine samples,^17^ a few recent studies proposed to longitudinally monitor serum steroid biomarkers.^6,12,16,18,19^ For instance, Salamin et al.^12^ monitored serum T, dihydrotestosterone (DHT), androstenedione (A4) as well as the ratio T/A4 and highlighted the relative stability of these compounds compared to the urine ones in addition to their greater sensitivity. Furthermore, serum steroids appear to be more stable over the menstrual cycle than urine compounds.^12,20^ As the first athletes were sanctioned for abnormal fluctuations in serum steroid levels^18^, the establishment and introduction of an upcoming ABP blood steroid module gained momentum.^21^

In this study, we aimed to monitor and comparie urine and serum steroid profiles over two consecutive OCP cycles among a physically active female population. We investigated the stability of serum parameters in the light of the upcoming ABP blood steroid module and assessed the potential confounding effects of the OCP cycle on individual profiles with the hypothesis that serum biomarkers would be more stable compared to urine variables and the OCP cycle would only elicit minor variations. In a second step we aimed to compare serum steroid concentrations in similar active women populations with and without OCP with the hypothesis that OCP users would present lower concentrations.

## Material and methods

### Study subjects

Fifteen healthy female subjects with regular withdrawal bleeding patterns, aged 23.2 ± 2.4 years volunteered to take part in this study and were monitored for 2 consecutive OCP cycles corresponding to 8 weeks. Recruitment was made by contacting students at the Institute of Sport Sciences of the University of Lausanne (ISSUL) and continued through a snowball sampling method.^22^ Subjects exercised for an average of 297 ± 100 min per week during the study whereas a minimum of 4 h of weekly physical activity was required to participate. All subjects had regular cycle length (28.5 ± 1.5 days) and were using oral contraceptive pill (more detailed anthropometric data are available eslewhere^23^). Intake of medication was recorded. Procedures and risks were fully explained to the subjects, and all of them gave their written consent to participate in the study. This study was approved by the local ethics committee (CER-VD, Lausanne, Switzerland, Agreement 2018-01019) and conducted in accordance with the Declaration of Helsinki. Secondly, data from Salamin et al.^12^ was used to compare OCP and non-OCP users. This study, approved by the local Ethical Committee of the Canton de Vaud in Switzerland (2018–02106, SNCTP000003264) and Swissmedic (2018DR1168) and also conducted in accordance with the Declaration of Helsinki, included fourteen healthy women aged 22-37 years (median 27.5 years) with regular menstruations and not using hormonal contraception.

### Study design

Urine and serum samples were collected once a week over 2 consecutive OCP cycles (8 weeks). Subjects started the study randomly regarding their OCP cycle and were asked to report to the laboratory each week during the two cycles at the same time of the day (to avoid putative circadian variations). Because of the COVID-19 pandemic, one subject had to self-isolate, and thus the eight samples were collected over two non-consecutive cycles in a 3-month period.

OCP cycle phases were subsequently assessed as weeks following the start of the withdrawal bleeding. This method was preferred to separating menstrual phases (follicular, ovulatory, and luteal phases) since OCP intake blunts the natural hormonal regulation of the menstrual cycle. However, due to its regular intake, the OCP does induce a cyclical bleeding which could be characterized as withdrawal bleeding. Therefore, in this paper we will be using the term OCP cycle instead of menstrual cycle.

### Sampling method

Participants reported to the lab having avoided any physical exercise in the 2 hours preceding the sampling and were asked to fill a disposable cup with their urine (Sarsted 100 mL sterile containers, Sarsted AG, Nümbrecht, Germany) and urine specific gravity was determined using a handheld refractometer (Pen-Urine SG, Atago, Fukui City, Japan). For serum sampling, blood was collected after 10 min seated in 9mL tubes (Sarstedt S-Monovette® Serum-Gel 9 mL, Sarstedt AG, Nümbrecht, Germany), considered equivalent from a clinical perspective to the BD Vacutainer® SST II Plus (EU ref 367955) proposed in the WADA blood collection guidelines.^24^ Venipuncture was realized with a 21G short manifold butterfly needle inserted into an antecubital vein (Sarstedt Safety-Multifly®, Sarstedt AG, Nümbrecht, Germany). Serum samples were centrifuged (Z326K Centrifuge, Hermle Labortechnik, Wehingen, Germany) for 10 minutes at 2500 rpm. They were finally aliquoted in three samples of 800 µl stored in a freezer at −20° C for later analyses.

### Urine analyses

Urine samples were prepared and analysed with gas chromatography coupled to tandem mass spectrometry (GC-MS/MS) according to the WADA technical document for the analysis of endogenous anabolic androgenic steroids (TD2021EAAS).^25^ The method is well described elsewhere.^26^ For the sample analyses from the OCP users cohort, the limit of quantification (LOQ) for T and E were 1 ng/mL; 20 ng/mL for A and Etio; and 2.5 ng/mL for 5αAdiol and 5ßAdiol. T, E, A, Etio, 5αAdiol and 5ßAdiol concentrations as well as specific gravity were entered into the Anti-Doping Administration and Management System (ADAMS) training environment according to the TD2021EEAS^25^ with approval by the WADA to compute individual longitudinal profiles.^5^

### Serum analyses

Serum levels of 3 different steroid hormones (T, A4 and DHT) were measured using a validated ultra-high performance liquid chromatography tandem mass spectrometry (UHPLC-MS/MS) method previously described.^19^ Calibration was performed using an 8-point linear calibration model (weighting 1/x) in spiked depleted serum for each compound, prepared freshly within each batch.

### Statistical analyses

Data are reported as means ± standard deviations (SD). Urine concentrations were adjusted for specific gravity. The variability of urinary and serum variables over time was calculated as coefficient of variations (CV%) from the mean of each individual’s CV over the 8 successive measures. Unpaired T test were used to assess the statistical significance for both CVs and the OCP and non-OCP conditions. In order to compare the OCP and non-OCP conditions in balanced datasets, data from Salamin et al.^12^ (non-OCP) were narrowed down to match the weekly sample collection in this study (OCP). Urinary and serum data were pooled in function of weeks following the withdrawal bleeding. P0 represents the week of the bleeding, P1 represents P0 plus 1 week, P2 represents P0 plus 2 weeks, and P3 represents P0 plus 3 weeks. The normality of the distributions was tested with the D’Agostino and Pearson test. Sphericity was not assumed, and the Geisser–Greenhouse correction was used. For normally distributed variables (A/Etio, serum T, A4, DHT and T/A4), mixed-effects analyses with Tuckey’s multiple comparisons test were performed. For the variables that were not normally distributed (A, Etio, 5αAdiol, 5ßAdiol and 5αAdiol/5ßAdiol), parameters were compared using a Friedman test with Dunn’s multiple comparisons. Pearson correlations were calculated for all variables. All data were analysed using a dedicated software (Prism, Version 8.4.2, GraphPad Software, La Jolla, California, USA).

## Results

### Urine steroids

When individual data were entered into ADAMS training environment software, no atypical passport finding (ATPF; e.g., value outside individual limit for the primary biomarker T/E) was reported. Nevertheless, one subject yielded a suspicious profile for one secondary biomarker with one low 5αAdiol/5ßAdiol value lying outside the individual calculated limits. In accordance with expected results from women urinary concentrations, T and/or E were below the LOQ for 74 samples which represents 62% of the total urine samples. Therefore, ratios dependent from T and E concentrations (5αAdiol/E and A/T) were not considered in statistical analyses due to the high number of missing values except from the T/E which is still reported when T and/or E are below LOQ.^25^

Individual urinary steroids values are reported as means, SD and CV in Table 1. The intra-individual variability over the 8-week study period was evaluated and the mean individual coefficient of variation remained comparably high for A (38%), Etio (41%), 5αAdiol (34%), 5ßAdiol (40%) and T/E (41%) while the A/Etio and 5αAdiol/5ßAdiol ratios respectively presented a significantly lesser variability with CVs of 20% and 25% (all p values < 0.05).

**Table 1.** Mean values, standard deviation (SD) and coefficient of variation (CV) for urinary testosterone (T), epitestosterone (E), androsterone (A), etiocholanolone (Etio), 5α-androstane-3α, 17β-diol (5αAdiol), 5β-androstane-3α,17β-diol (5βAdiol), T/E, A/Etio and 5αAdiol/5βAdiol for the 8 consecutive weeks of measurement.

| | T<br>(ng/mL)<br>(n=59) | E<br>(ng/mL)<br>(n=48) | A<br>(ng/mL)<br>(n=119) | Etio<br>(ng/mL)<br>(n=119) | 5αAdiol<br>(ng/mL)<br>(n=119) | 5βAdiol<br>(ng/mL)<br>(n=119) | T/E<br>(n=119) | A/Etio<br>(n=119) | 5αAdiol/<br>5βAdiol<br>(n=119) |
| --- | --- | --- | --- | --- | --- | --- | --- | --- | --- |
| **Mean** | 7.1 | 5.0 | 1412 | 1802 | 17.0 | 60.7 | 2.05 | 0.9 | 0.4 |
| **SD** | 2.5 | 1.7 | 572 | 764 | 6.1 | 24.7 | 0.80 | 0.2 | 0.1 |
| **CV (%)** | 31.3 | 32.1 | 38.1 | 40.8 | 34.2 | 40.0 | 41.4 | 19.7 | 25.2 |

When considering data pooled in function of weeks following the withdrawal bleeding, significant statistical differences occurred for 5ßAdiol, T/E and A/Etio. Mean 5ßAdiol decreased by 47% (p<0.05) from P1 (78.1 ± 98.9 ng/mL) to P2 (53.0 ± 63.23 ng/mL). The mean ratio T/E increased significantly at P2 (2.4 ± 1.3) compared to P0 (1.7 ± 0.9; p < 0.05) and P1 (1.8 ± 1.0; p < 0.05) and at P3 (2.3 ± 1.5) compared to P0 (p < 0.05) while A/Etio increased significantly by more than 17% (p<0.05) at P2 and P3 compared to P0 (0.79 ± 0.30 vs. 0.96 ± 0.39 (p < 0.05) and 0.93 ± 0.30 (p < 0.05)) and P1 (0.79 ± 0.34 vs. 0.96 ± 0.39 (p<0.01) and 0.93 ± 0.30 (p<0.05)). The median, quartile, maximal and minimal values in the weeks following the bleeding for A, Etio, 5αAdiol, 5ßAdiol, A/Etio and 5αAdiol/5ßAdiol are illustrated in Figure 1.

**Figure 1.**
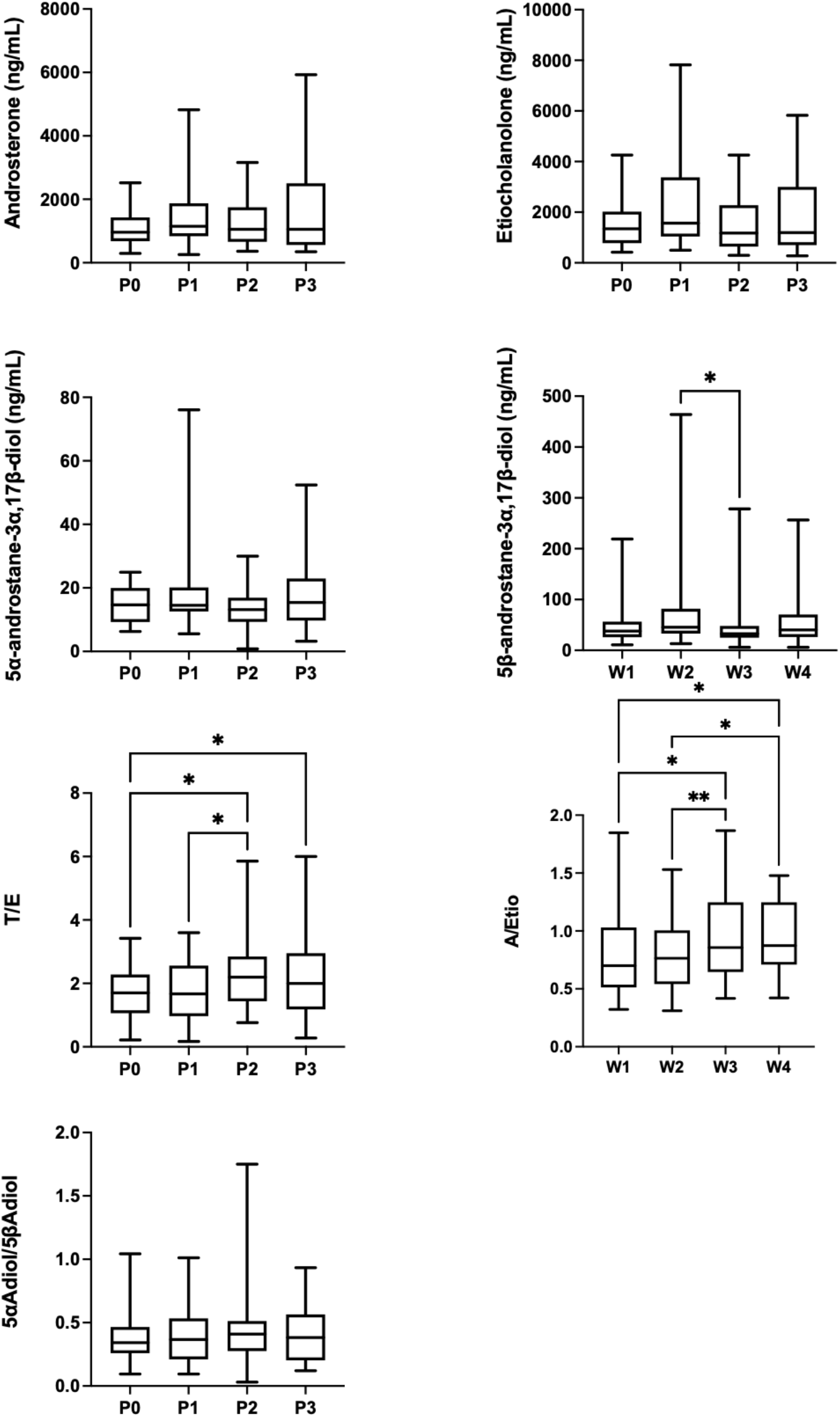
Urinary androsterone (A), etiocholanolone (Etio), 5α-androstane-3α, 17β-diol (5αAdiol), 5β-androstane-3α,17β-diol (5βAdiol), T/E, A/Etio and 5αAdiol/5βAdiol in the phases following the week of the withdrawal bleeding. P0 represents the week of the withdrawal bleeding, P1 represents P0 plus 1 week, P2 represents P0 plus 2 weeks, and P3 represents P0 plus 3 weeks. Error bars represent minimum to maximum. ** p < 0.01 and * p < 0.05 for the difference between phases.

### Serum steroids

Individual serum steroids values are reported as means, SD with CV in Table 2. The intra-individual variability was lower for serum steroids than urines (p < 0.001). Mean testosterone concentrations in urine and serum samples collected during the 8 successive weeks are presented comparatively in Figure 2. CVs for T and A4 were respectively 24.4% and 22.8% while DHT had a significantly lesser variability than T (18.6%; p < 0.05) despite a higher number of values below LOQ (20%). When combining T and A4 into the T/A4 ratio, variability further decreased to 15.6% which is significantly lower to the CVs of T, A4 and DHT (all p values < 0.001). It is also noticeable that no significant correlation was observed between urinary and serum T for any subject nor between any variables considered in this study.

**Table 2.**
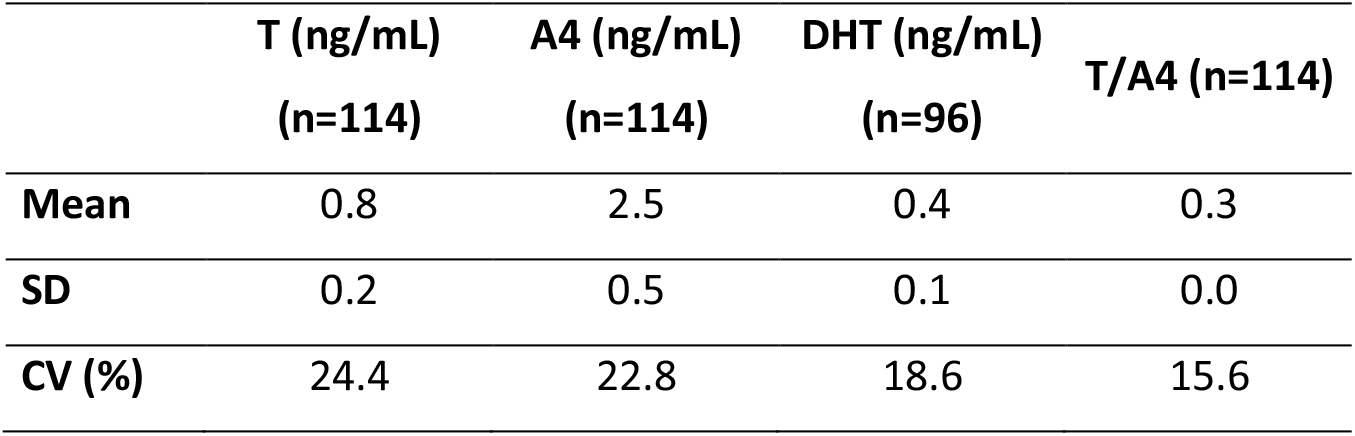
Mean values, standard deviation (SD) and coefficient of variation (CV) for serum testosterone (T), androstenedione (A4), dihydrotestosterone (DHT) and T/A4 for the 8 consecutive weeks of measurement.

**Figure 2.**
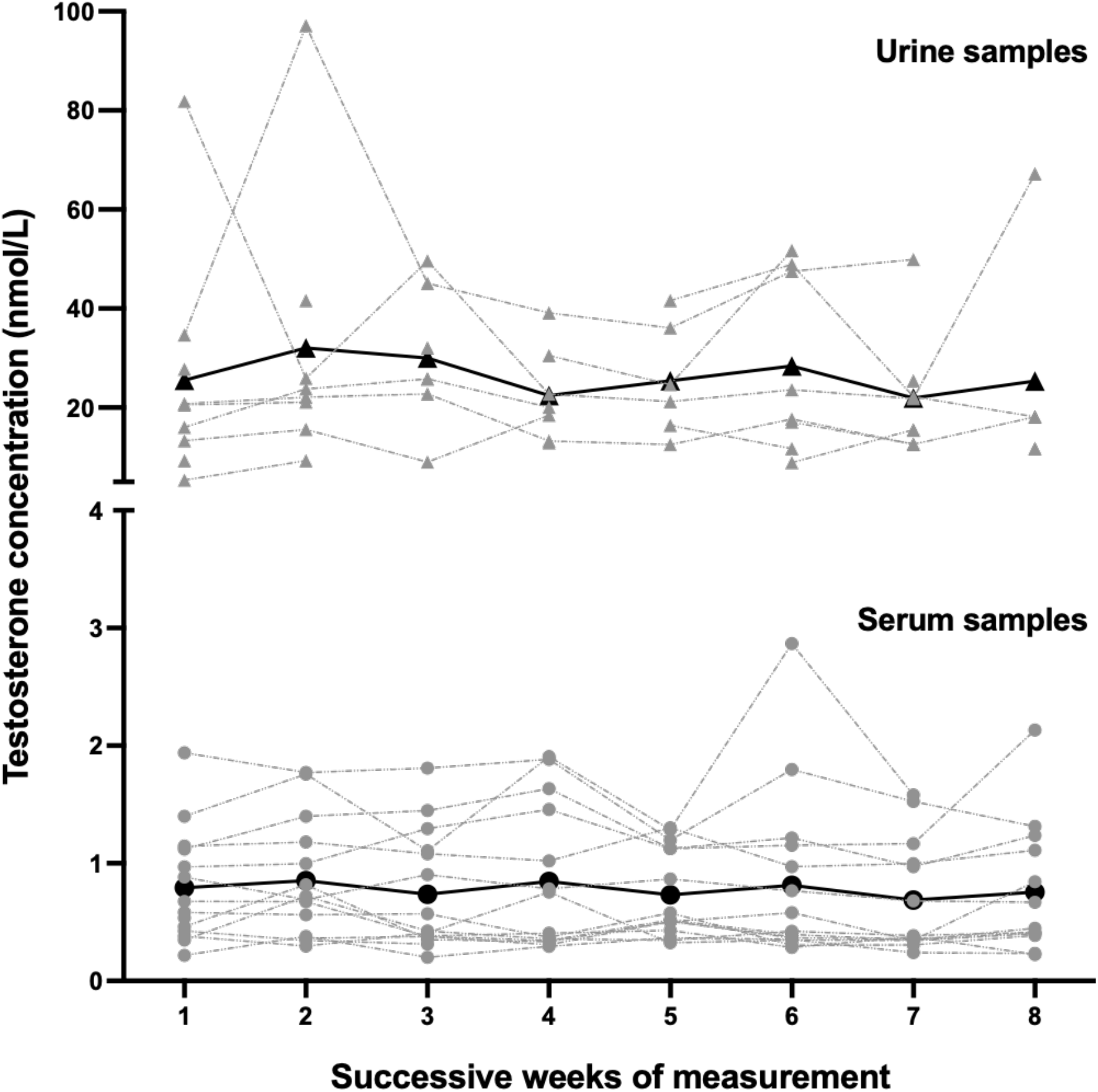
Urinary testosterone (nmol/L) (triangles; n=59) and steroid testosterone (nmol/L) (circles; n=114) for every subject during the 8 successive weeks of measurement. Black lines represent mean values.

When pooling the data in function of weeks following the withdrawal bleeding, T and T/A4 were significantly higher at P0 (0.98 ± 0.63 nmol/L; 0.34 ± 0.12) compared to P1 (0.73 ± 0.46 nmol/L; p < 0.01; 0.28 ± 0.09; p < 0.001; respectively), P2 (0.70 ± 0.50 nmol/L; p < 0.001; 0.28 ± 0.10; p < 0.001; respectively) and P3 (0.71 ± 0.48 nmol/L; p < 0.01; 0.28 ± 0.11; p < 0.001; respectively) whereas the only significant difference for A4 is at P0 (2.78 ± 1.07 nmol/L) compared to P2 (2.33 ± 1.03 nmol/L; p < 0.05). DHT showed no significant variation over the OCP cycle. The median, quartile, maximal and minimal values in the weeks following the bleeding for T, A4, DHT and T/A4 are illustrated in Figure 3.

**Figure 3.**
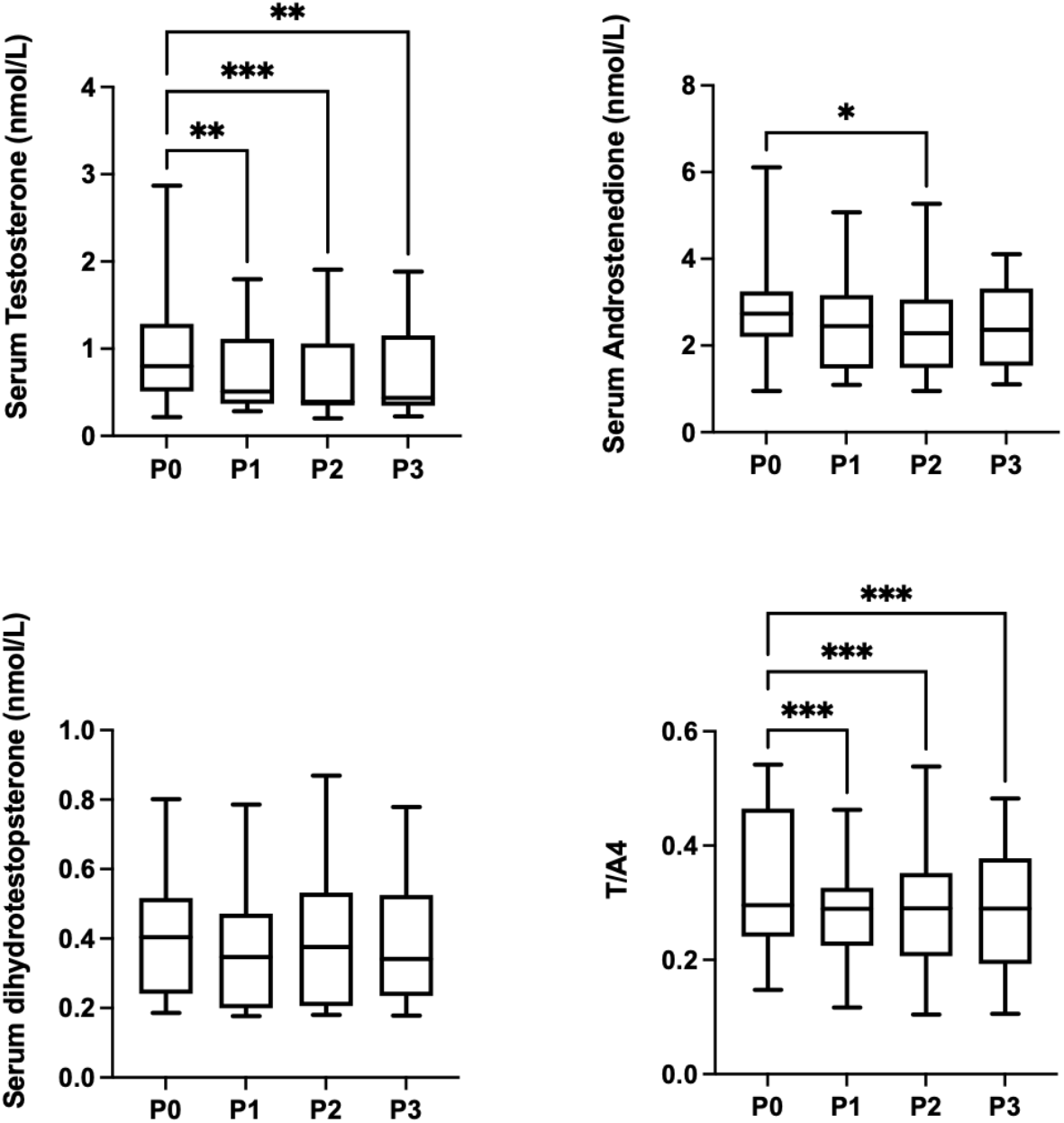
Serum testosterone (T), androstenedione (A4), dihydrotestosterone (DHT) and T/A4 in the phases following the week of the withdrawal bleeding. P0 represents the week of the withdrawal bleeding, P1 represents P0 plus 1 week, P2 represents P0 plus 2 weeks, and P3 represents P0 plus 3 weeks. Error bars represent minimum to maximum. *** p < 0.001, ** p < 0.01 and * p < 0.05 for the difference between phases.

### OCP use

When the non-OCP and OCP cohorts were compared, some biomarkers concentrations were significantly lower for OCP users in both urine and serum. In urine, concentrations were significantly lower with OCP intake for E (15.1 ± 8.4 ng/mL vs. 5.1 ± 2.6, p < 0.001), A (2399 ± 1448 ng/mL vs. 1417 ± 1076 ng/mL, p < 0.001), Etio (2758 ± 1797 ng/mL vs. 1811 ± 1349 ng/mL, p < 0.001) and 5αAdiol (28.5 ± 17.4 ng/mL vs. 17.0 ± 10.6 ng/mL, p < 0.001). Concomitantly to those differences, T/E (0.55 ± 0.42 vs. 2.06 ± 1.21, p < 0.001) and 5αAdiol/E (2.2 ± 1.1 vs. 4.1 ± 4.9, p < 0.001) ratios present significantly higher values for OCP users while A/T (815 ± 1324 vs. 275 ± 132, p < 0.001) is significantly lower. T (7.0 ± 5.3 ng/mL vs. 7.6 ± 5.1 ng/mL), 5ßAdiol (68.8 ± 46.0 ng/mL vs. 61.1 ± 69.6 ng/mL), A/Etio (0.9 ± 0.3 vs. 0.9 ± 0.3) and 5αAdiol/5ßAdiol (0.5 ± 0.2 vs. 0.4 ± 0.3) showed no statistically significant difference. The biomarkers data distribution is illustrated in Figure 4. In serum, concentrations of all biomarkers (T: 1.1 ± 0.4 nmol/L vs. 0.8 ± 0.5 nmol/L, p < 0.001; A4: 5.2 ± 1.7 nmol/L vs. 2.5 ± 1.0 nmol/L, p < 0.001; and DHT: 0.6 ± 0.4 nmol/L vs. 0.4 ± 0.2 nmol/L, p < 0.001) were significantly lower in OCP users while the T/A4 ratio was concomitantly and significantly higher (0.22 ± 0.05 vs. 0.30 ± 0.10, p < 0.001). The biomarkers data distribution is illustrated in Figure 5. No significant correlation was observed neither between oestrogens and progesterone nor in the other analysed biomarkers.

**Figure 4.**
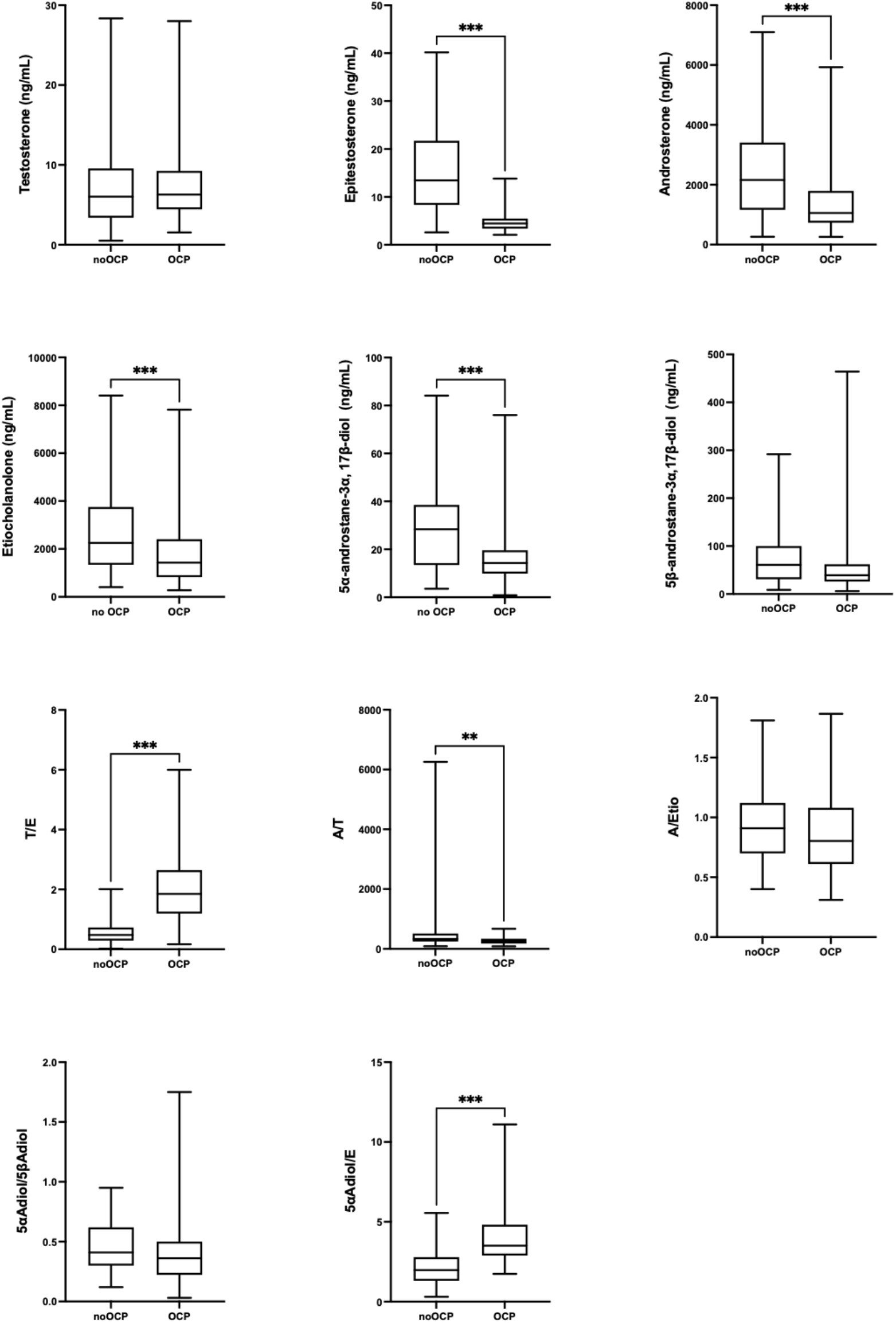
Urinary androsterone (A), etiocholanolone (Etio), 5α-androstane-3α, 17β-diol (5αAdiol), 5β-androstane-3α,17β-diol (5βAdiol), A/Etio and 5αAdiol/5βAdiol for both non-OCP and OCP conditions. Error bars represent minimum to maximum. *** p < 0.001, ** p < 0.01 for the difference between conditions.

**Figure 5.**
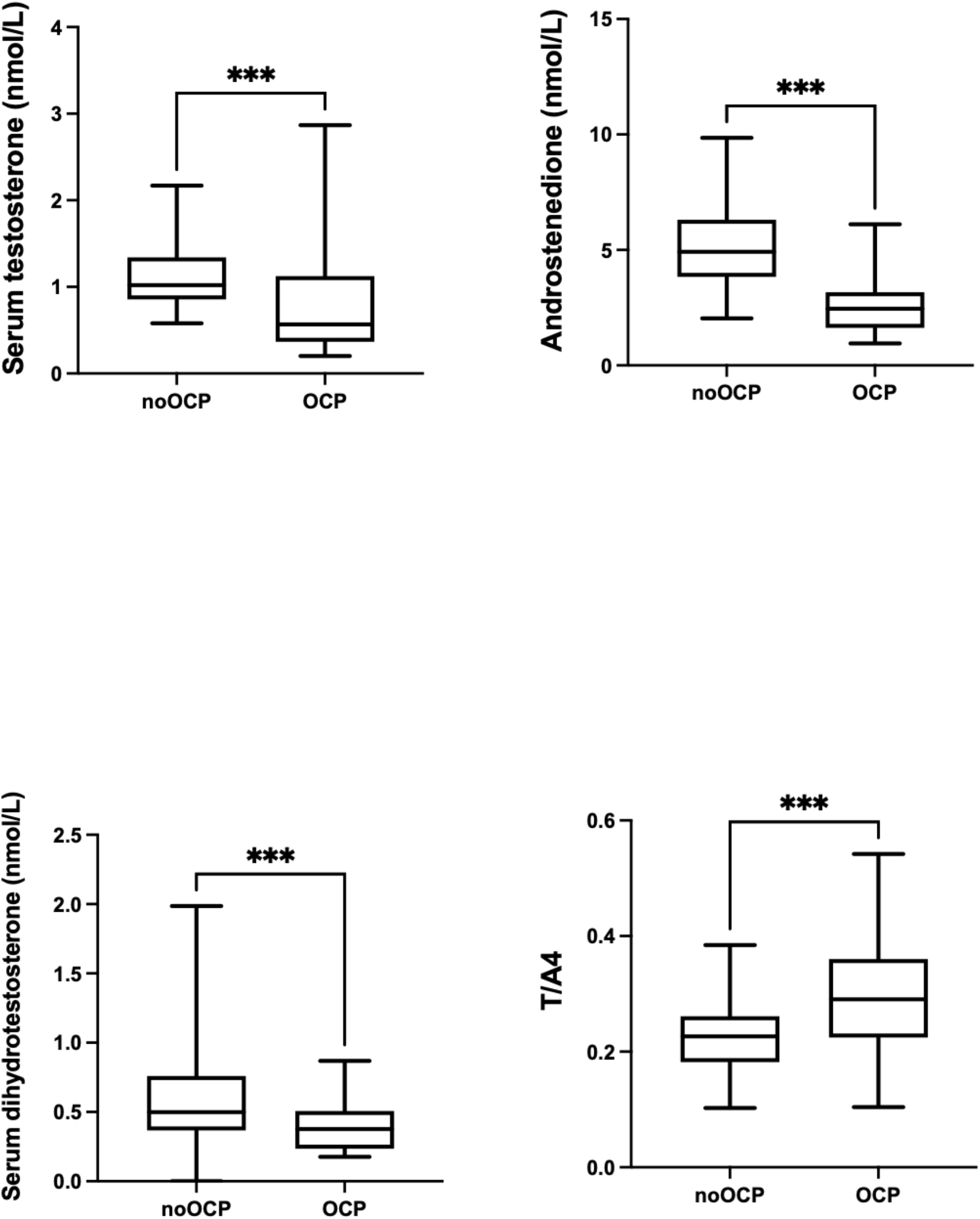
Serum testosterone (T), androstenedione (A4), dihydrotestosterone (DHT) and T/A4 for both non-OCP and OCP conditions. Error bars represent minimum to maximum. *** p < 0.001 for the difference between conditions.

## Discussion

In the present study, the variability of the urine and blood steroid profile was assessed and compared over 8 weeks corresponding to two OCP cycles in 15 healthy women. This is the first time that urinary and serum steroid levels for both OCP and non OCP users are compared in similar healthy active women. The main finding of this study is that T and/or E were below the LOQ (1 ng/mL) for 62% of the urine samples (n=74 out of 119). This is line with a recent study from Schulze et al. (2021) similarly reporting a considerable number of missing values for the T and/or E in urine (6 subjects out of 17). Urine steroids levels were however in agreement with low reference ranges for women reported in the literature. ^6,8–10^ Due to a majority of values below the LOQ T and E were not considered in statistical analyses but would have indisputably impacted the respective mean values that can be statistically reported and should be considered to avoid any misinterpretation of the actual concentrations. Since values below LOQ may impair the calculation of upper and lower individual limits in the ABP adaptive model, individual profiles of women with low steroidal concentrations might prove to be more difficult to interpret. For instance, all urinary and serum sequences of data for both urine and serum T for all the 15 subjects in Figure 2 highlight missing values in the urinary samples due to low concentrations. In contrast, full serum sequences can be obtained in all subjects which highlights the ability of serum steroids to produce more robust data for a pertinent interpretation of individual profiles. However, the deletion polymorphism of UGT2B17 enzyme (known to induce very low T excretion concentrations) was not controlled in our study cohort. It is however unlikely that this polymorphism was present in high number in the study cohort.

In contrast, serum steroids concentrations presented considerably lower intra-individual variations compared to urine with concentrations for our study cohort in accordance with results reported earlier.^6,16,27,28^ For illustration, Subject D had a mean CV of 59% for all urine biomarkers whereas the overall variability for serum biomarkers variability was of 25% (see Tables 1 and 2). Higher intra-individual variability in urine biomarkers and ratios can prove to be detrimental to the ABP sensitivity with its impact on the generation of upper and lower individual limits.

Serum may hence definitely represent a complementary matrix to urine in the context of the ABP. First, analytical methods can accurately detect lower absolute concentrations of steroids which can prove to be critical in maintaining a high sensitivity score in the ABP, especially for women with low steroidal levels. Steroid monitoring in serum allowed better detection of AAS administration than traditional urine follow-up in several recent studies.^12,16,18^ Furthermore, serum biomarkers are more stable than urine’s and would help increase the ABP sensitivity. It has recently been suggested by Salamin et al.^12^ that, in women, the combination of T and A4 into a ratio improves the stability and the detection capacity of longitudinal profiles. This is well illustrated in our study with T/A4 presenting the lowest variability of all monitored variables. Therefore, including serum biomarkers in the ABP can be a useful tool to target testing and to increase indirect detection of EAAS abuse.^17^

To assess putative recurring patterns during the OCP cycle in urinal and serum steroids, data was pooled in function of weeks following the withdrawal bleeding. The significant increase in T/E at P2 is related to a decrease in E, which is in line with the reported large inter-individual difference and impact of OCP on hormonal metabolism.^13–15^ The significant increase in A/Etio in the luteal phase (Figure 1) contrasts with a previous study on women without OCP^11^ supporting therefore an effect from OCP intake on subsequent hormonal concentrations in urine. The other urinary biomarkers did not vary significantly (except from a small significant increase at P1 compared to P2 for 5ßAdiol), in line with the literature.^11,29,30^ In serum, the significant decrease of T and T/A4 during the 3 weeks following the bleeding (Figure 3) is contradictory again with the stable T reported earlier but in women without OCP.^11^ Again, this discrepancy may be explained by the hormonal intake during the OCP cycle since the significant increase of T and T/A4 occurred during the OCP placebo phase. This latter finding is paramount while WADA is implementing an ABP blood steroid module with the OCP cycle to be considered as an important confounding factor in females.^21^ For instance, hormones contained in OCP preparations may blunt steroid metabolism due to a negative feedback mechanism.^31^ However, most OCP regimens include a placebo phase in which steroid metabolism tend to revert endogenous steroids to natural levels. This increase remains modest and is arguably unlikely to have a significant effect on longitudinal monitoring and yield atypical findings in the ABP adaptative model.

When urinary and serum steroid concentrations of this study were compared to the data from Salamin et al.^12^, OCP users had significantly lower levels than non-users with the most impacted biomarkers being E in urine and A4 in serum (Figures 4 and 5). Data dispersion for E in OCP users is remarkably low and highlights the putative effect of OCP (Figure 4). This result is well known and in line with the literature reporting lower urinary steroid levels with OCP.^13–15^ However, T and E values for OCP users are certainly largely overestimated as this cohort is characterized by a high number of values below the LOQ that could not be included. Regarding serum biomarkers, while decreased levels of T were found in clinical contexts, data on other biomarkers of the upcoming blood steroid profile are scarce.^32–34^ Our results show that A4 is the most impacted variable which concomitantly reverses the difference for the T/A4 ratio (Figure 5) whereas, interestingly, a recent work from Knutsson et al.^35^ found that T/A4 remained unaffected by the intake of OCP as both T and A4 varied to the same extent. Here, even though T/A4 is the marker presenting the smaller variation (+24%), it increases significantly due to the larger decrease of A4 concentrations (−52%) compared to T (−31%). Moreover, A4 and DHT present smaller data dispersion which underlines an uniformizing effect of OCP. The comparison in variability between urinary and serum steroidal biomarkers presented here can prove to be very helpful for future expert reviewing the upcoming blood steroid profile. While Salamin et al.^12^ noted significant correlations between both estrogens and progesterone and E in a non-OCP cohort, no such correlation was observed in OCP users. This is not surprising as OCP blunts those hormones metabolism and maintain low concentration level.

## Conclusion

In this study, urine and serum steroids were monitored over two consecutive OCP cycles in physically active women. The very low steroid concentrations in female athletes were outlined by a considerable number of missing values for urinary steroids that may negatively impact the interpretation of individual profiles in an ABP context. This study therefore supports the incorporation of serum as a new matrix to monitor steroids, as both sensitivity and selectivity of the ABP could be improved by the greater stability of the steroid biomarkers in serum. When comparing both urinary and blood steroid concentrations between non-OCP and OCP users, most biomarkers showed significatively lower levels with OCP intake. In conclusion, our results suggest that the OCP cycle has an influence on some parameters in both urine and serum with significant variations in urine’s A/Etio, serum’s T and T/A4. The use of OCP in female athletes must be considered as a confounding factor of the forthcoming blood steroid module of the ABP. A thorough investigation and review of all potential confounders when investigating serum and urinary steroidal biomarkers may represent the next step in the direct and indirect detection of AAS abuse in female athletes.

## Authors’ contributions

RF designed the study and obtained funding. BM, BK, JJS and RF contributed to data collection. OS, RN, and TK performed and reviewed the serum analyses. LI, FM, and FB performed and reviewed the urinary analyses. BM drafted the first version of the manuscript; FB, OS, JJS and RF revised it critically. All authors read and approved the final version of the manuscript.

## Acknowledgements

The authors wish to acknowledge WADA’s Science Department for the financial support of this study and all the participants for their participation.

